# IL-34 deficiency impairs FOXP3^+^ Treg function and increases susceptibility to autoimmunity

**DOI:** 10.1101/2022.02.04.479184

**Authors:** Antoine Freuchet, Apolline Salama, Séverine Bézie, Laurent Tesson, Séverine Rémy, Romain Humeau, Hadrien Règue, Léa Flippe, Pärt Peterson, Nadège Vimond, Claire Usal, Séverine Ménoret, Jean-Marie Heslan, Franck Duteille, Frédéric Blanchard, Magali Giral, Marco Colonna, Ignacio Anegon, Carole Guillonneau

**Affiliations:** Nantes Université, Inserm, Center for Research in Transplantation et Translational Immunologie, UMR 1064, ITUN, F-44000 Nantes, France; Institute of Biomedecine and Translational Medecine, University of Tartu, Tartu, Estonia; Nantes Université, CHU Nantes, Inserm, CNRS, SFR Santé, Inserm UMS 016, CNRS UMS 3556, F-44000 Nantes, France; Chirurgie Plastique Reconstructrice et Esthétique, CHU Nantes, Nantes, France; INSERM UMR1238, Bone Sarcoma and remodeling of calcified tissues, Nantes University, 44093 Nantes, France; Department of Pathology and Immunology, Washington University School of Medecine, St. Louis, MO 63110, USA

**Keywords:** IL-34, Treg, tolerance, autoimmunity, immunotherapy, knockout

## Abstract

Immune homeostasis requires fully functional Tregs with a stable phenotype to control autoimmunity. Although IL-34 is a cytokine first described as mainly involved in monocyte cell survival and differentiation, we recently described its expression by CD8+ Tregs in a rat model of transplantation tolerance and by activated FOXP3+ CD4+ and CD8+ Tregs in human healthy individuals. However, its role in autoimmunity and potential in human diseases remain to be determined. Here we report that the absence of expression of IL-34 in *Il34*^-/-^ rodents leads to an unstable phenotype, with production of multiple auto-antibodies, exacerbated under inflammatory conditions with increased susceptibility to DSS- and TNBS-colitis in deficient animals. Moreover, we strikingly revealed the inability of *Il34*^-/-^ CD4+ Tregs to protect *Il2rg*^-/-^ rats from a wasting disease induced by transfer of pathogenic cells, in contrast to *Il34*^+/+^ CD4+ Tregs. We also showed that IL-34 treatment delayed EAE in mice as well as GVHD and human skin allograft rejection in immune humanized immunodeficient NSG mice. Finally, we show that presence of IL-34 in the serum is associated with a longer rejection-free period in kidney transplanted patients. Altogether, our data emphasize on the crucial necessity of IL-34 for immune homeostasis and for CD4+ Tregs suppressive function. Our data also shows the therapeutic potential of IL-34 in human transplantation and auto-immunity.

## Introduction

IL-34 is a homodimeric cytokine binding three distinct receptors or co-receptors, namely CSF-1R (CD115), CD138 (syndecan-1) and PTPζ (1, 2). CSF-1 is another ligand of CD115 but binds with a lower affinity than IL-34, and they both induce survival and differentiation of monocytes toward type 2 “regulatory” macrophages (3, 4). However, IL-34 and CSF-1 spatio-temporal expressions are mainly distinct and CSF-1 does not bind to CD138 and PTPζ (5). Thus, they have been shown to exert also non-overlapping roles (5).

We have recently demonstrated that IL-34, but not CSF-1, is significantly overexpressed in antigen-specific CD8+ Tregs from long-term tolerant transplanted rats (treated with CD40Ig, a chimeric molecule blocking the CD40-CD40L pathway) vs. CD8+ Tregs from naive rats (6–8). Further analysis showed that IL-34 expression in human T cells is restricted to, not only activated FOXP3+ CD8+ Tregs, but also to FOXP3+ CD4+ Tregs with about 50% of the FOXP3+ Tregs expressing IL-34 (6, 9). We also showed that treatment using an AAV encoding IL-34 alone or together with a short sub-optimal dose of rapamycin efficiently delayed or inhibited allograft rejection *in vivo* in rats through differentiation of tolerogenic macrophages responsible, in turn, for induction of CD4+ and CD8+ Tregs. In this model, the CD4+ and CD8+ Tregs were responsible for the long-term tolerance induction effect of the treatment. *In vitro*, we showed that IL-34 inhibited both rats and humans effector CD4+CD25^-^ T cells proliferation and induced human Foxp3+ CD4+ and CD8+ Tregs expansion (6, 10). In addition, a suppressive role for IL-34 was also demonstrated by others (11) in rodent liver transplantation models.

While *Il34*^*LacZ*/*LacZ*^ mice have been studied and no spontaneous autoimmunity or inflammatory diseases were reported (12), we speculated that it could be due to lack of an inflammatory/autoimmune stimulus. Indeed, in human, increased IL-34 serum levels have been associated to inflammatory diseases (1, 2), however IL-34 role was not properly described and mostly circumstantial. Thus, the contribution of IL-34 in immune homeostasis, Treg function, as well as its role in autoimmunity and human transplant rejection remains unclear and need to be further addressed. Herein, we generated *Il34*^-/-^ rats and using both *Il34*^-/-^ rats and mice, we investigated their phenotype under inflammatory conditions. Using *Il34*^-/-^ rats, we further analyzed the impact of the absence of expression of IL-34 for CD4+ Tregs suppressive function. We investigated the potential of IL-34 in human disease to prevent xenogeneic GVHD and human skin allograft rejection in immune humanized immunodeficient NSG mice. Finally, taking advantage of a biocollection, we investigated the correlation between presence of IL-34 in the serum and kidney transplant rejection.

Altogether, our study emphasizes the crucial role of IL-34 for immune homeostasis and for CD4+ Tregs suppressive function and the therapeutic potential of IL-34 in human transplantation and auto-immunity.

## Results

### At homeostasis, IL-34 participates in microglia development and regulates cytokines, enzymes and auto-antibody production

To gain further knowledge on the immunoregulatory role of IL-34, we generated an *Il34* deficient rat model using CRISPR/Cas9 targeting exon 3 of the rat *Il34* gene. The Non-Homologous End Joining (NHEJ) repair mechanism inserted a C base, leading to a STOP codon apparition (**Figure 1A and Supplemental Figure 1A**). The founder animal was mated with an *Il34*^+/+^ animal, the mutation was transmitted to the progeny and homozygous *Il34*^-/-^ animals were generated as confirmed by heteroduplex mobility assay on an automated microfluidic chip capillary electrophoresis system and sequencing (13, 14). In absence of available anti-rat IL-34 antibody, we confirmed the knock-out using quantitative RT-PCR and observed a major decrease of *Il34* mRNA expression in *Il34*^-/-^ animals compared to littermates *Il34*^+/+^ animals (**Figure 1B**). Residual mRNA detected was due to mRNA being produced with an immature STOP codon before rapid degradation. The *Il34*^-/-^ animals looked visually healthy at steady state with normal appearance (**Supplemental Figure 1B**), growth (**Supplemental Figure 1C**) and bone mass (**Supplemental Figure 1D, E**). However, in accordance with the role of IL-34 reported for microglia development and maintenance in the mouse (5), brain analysis showed decreased microglia (CD11b/c+ cells) in the hippocampus, but not the cerebellum of the *Il34*^-/-^ animals (**Figure 1C, D and Supplemental Figure 1F**). Sera analysis showed an increase of ALT and ALT/LDH enzyme levels ratio and a trend for AST and alkaline phosphatase (**Figure 1E**), suggesting mild liver injury. Interestingly, we also observed a strong increase of auto-antibodies production against IFN-α1, −α2, −α4, -α7, -α11 and anti-IL-22 in *Il34*^-/-^ rats (**Figure 1F**) (no evidence for anti-IL-17A/F Ab), increased inflammatory chemokine MIP-2 and decreased eosinophil chemotactic proteins (Eotaxin), as well as regulatory TGF-β3 (**Figure 1G**) (no changes for TGFβ1/2, IL-1α /β, -2, -4, -5, -6, -10, -12p70, -13, -17A, G-CSF, GM-CSF, TNFα, IFNγ, GROα, MCP-1, -3, MIP-1α, Rantes and IP-10; **Supplemental Figure 2A**), altogether suggesting auto-inflammation and an important role for IL-34 in auto-antibody development and cytokine production. Serum levels of immunoglobulin isotypes were normal and antibodies directed against dsDNA were similar to *Il34*^+/+^ rats (**Supplemental Figure 2B-C**). Histological analysis of organs did not show evidence of tissue lesions or inflammatory infiltrates (**Supplemental Figure 2D**). We investigated whether, as compensatory mechanisms that could mitigate the auto-immune symptoms, CSF-1 was upregulated and indeed found significantly higher levels of CSF-1 in the sera of the *Il34*^-/-^ animals (>6 months old) (**Figure 1H**), suggesting a negative regulation of CSF-1 by IL-34.

**Figure 1.**
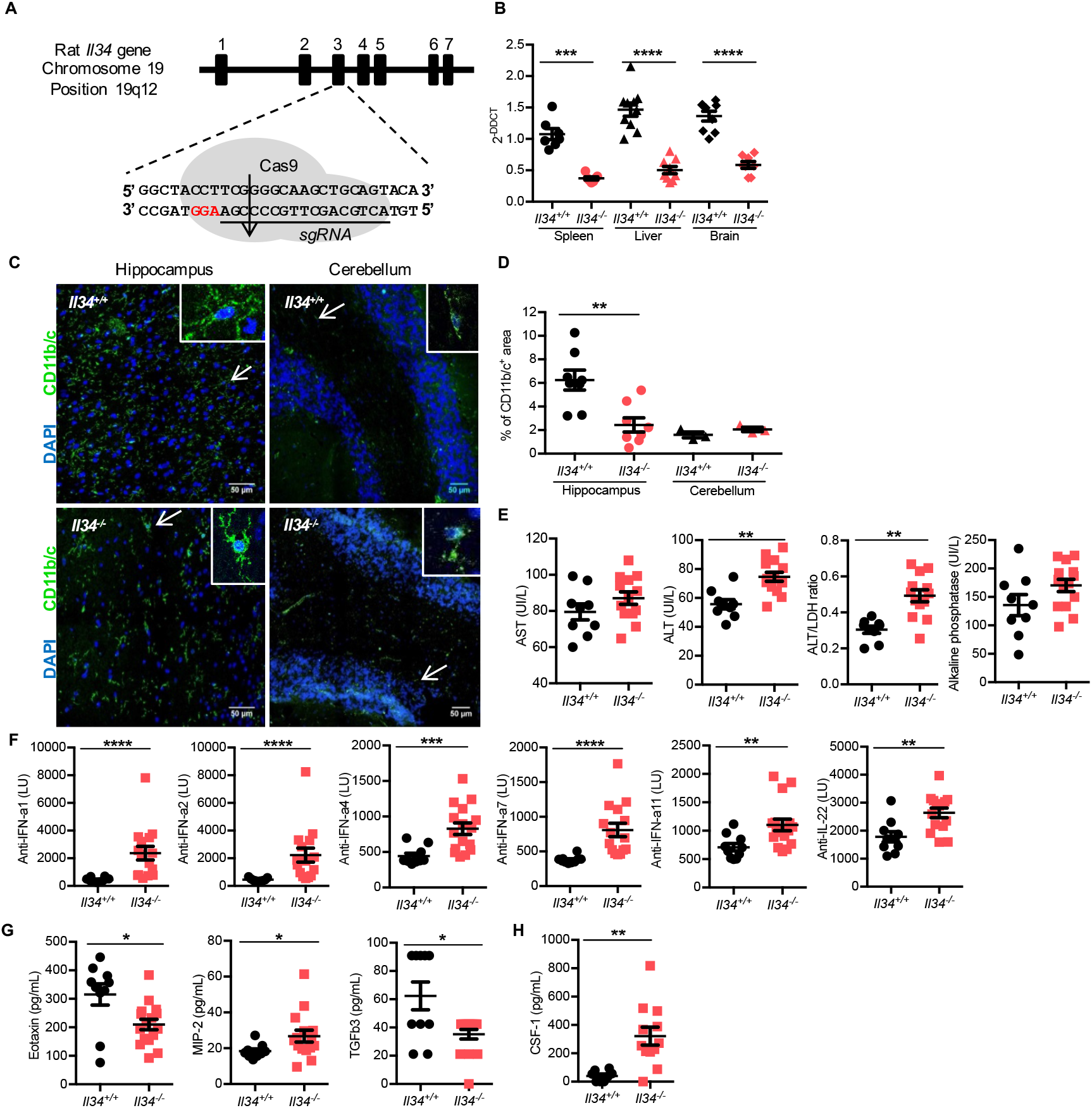
Generation of *Il34*^-/-^ rats by CRISPR/Cas9 and characterization. **(A)** Schematic representing the Cas9/sgRNA targeting the exon 3 of the *Il34* gene inducing a genomic DNA cut (arrow). PAM sequence is in red. **(B)** *Il34* mRNA expression was assessed by quantitative real-time PCR in spleen, liver and brain of *Il34*^+/+^ (n=7-10) and *Il34*^-/-^ rats (n=7-9). Results are normalized to *Hprt* and expressed as 2^-ΔΔCT^ ± SEM. **(C)** Confocal microscopy was performed on *Il34*^+/+^ and *Il34*^-/-^ rats frozen brains stained with an antibody directed against CD11b/c (green) to identify microglia and DAPI (blue). Arrows indicate a representative stained cell. Original magnification, x800. Scale bar 50 μm. **(D)** Quantification of positive CD11b/c staining areas in *Il34*^+/+^ and *Il34*^-/-^ rats’ hippocampus (*Il34*^+/+^, n=8; *Il34*^-/-^, n=8) and cerebellum (*Il34*^+/+^, n=3; *Il34*^-/-^, n=3) slices. Each dot represents an individual animal and results are expressed as mean ± SEM. **(E)** Plasma from 4 months old *Il34*^+/+^ (n=9) and *Il34*^*-/*-^ (n=14) rats was used to quantify AST, ALT, ALT/LDH ratio and alkaline phosphatase. **(F)** Auto-antibodies against interferon (IFN)-α 1, α 2, α 4, α 7, α 11 and IL-22 were assessed in sera by LIPS assay (>6 months old *Il34*^+/+^ rat, n=10 and *Il34*^-/-^ rat, n=16). **(G)** Eotaxin, TGF-β3 and MIP-2 protein levels were quantified by Luminex assay in the sera of >6 months old *Il34*^+/+^ (n=10) and *Il34*^-/-^ (n=16) rats. **(H)** CSF-1 protein was quantified by ELISA in the sera of >6 months old *Il34*^+/+^ (n=8) versus *Il34*^-/-^ rats (n=12). Results are expressed as mean ± SEM. Mann Whitney *U* test, * *p*<0.05, ** *p*<0.01, *** *p*<0.001, **** *p*<0.0001.

Altogether, these results demonstrate a role for IL-34 in rat microglia development and a critical role for regulating auto-antibody development and cytokine production.

### In rat, IL-34 deficiency leads to decreased number of CD8+ T cells and an increased susceptibility to colitis

We then further analyzed the impact of the deficiency on immune cells in mice and rats. While absolute numbers of cells were slightly decreased in thymus, spleen and blood of *Il34*^-/-^ rats (**Figure 2A**), thymic subpopulations, B cell subsets and myeloid cell subsets in spleen were not modified (**Supplemental Figures 3A-C**). In contrast, we observed a significant decrease of CD8+ T cells from spleen and blood in *Il34*^-/-^ rats vs *Il34*^+/+^ rats, but not for CD4+ T cells (**Figure 2B**). In addition, further analysis of lymphocyte function revealed a higher proliferation capacity of CD8+CD45RC^high^ (effector) T cells (10, 15, 16), but not of CD4+CD25^-^ effector T cells, in response to a polyclonal stimulation (**Figure 2C**), altogether suggesting a role for IL-34 in CD8+ T cell number and proliferative capacity.

**Figure 2.**
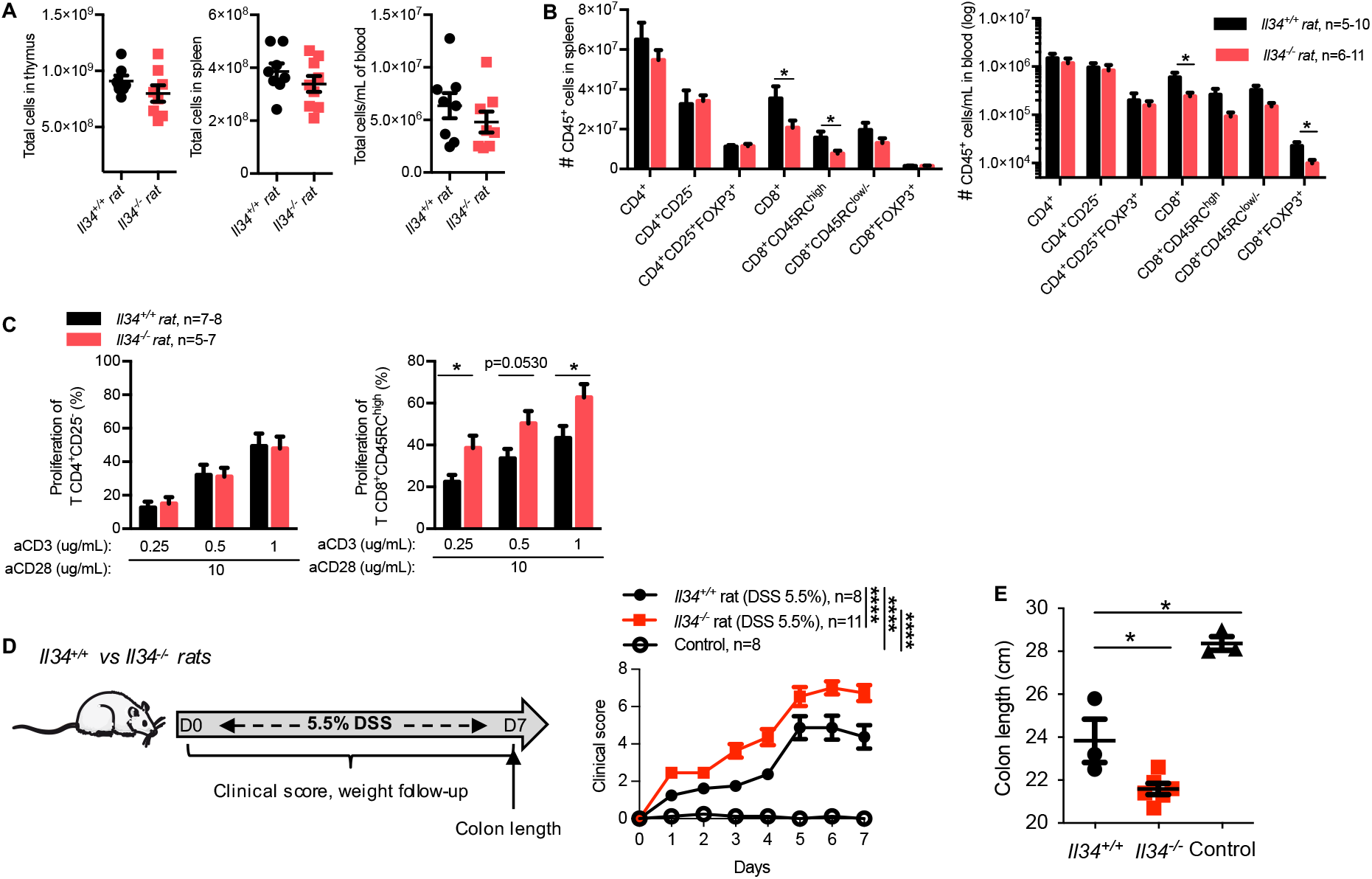
IL-34 global deficiency increases CD8+ T cells proliferation capacity and colitis severity. **(A)** Absolute numbers of cells were analyzed in thymus (2 months old), spleen and blood (>10 months old) of *Il34*^+/+^ (n=5-10) and *Il34*^-/-^ (n=6-11) rats. **(B)** Absolute numbers of T cell subsets were analyzed in spleen (left) and blood (right) of >10 months olds *Il34*^+/+^ (n=5-10) and *Il34*^-/-^ (n=6-11) rats using markers described in *Supplemental Methods*. **(C)** Effector TCRαβ+CD4+CD25^-^ (CD4+ Teffs) and TCRαβ+CD4^-^CD45RC^high^ (CD8+ Teffs) from *Il34*^+/+^ (n=7-8) or *Il34*^-/-^ (n=5-7) rats were sorted and tested for proliferation capacity over 2 days culture with increasing concentration of anti-CD3 (0.25-0.5-1 ug/mL) and anti-CD28 (10 ug/mL) mAbs. Results are represented as mean ± SEM. **(D**) *Left panel*. Schematic of the DSS model: 5.5% of DSS in drinking water was given for 7 days to 9-weeks old male *Il34*^+/+^ (n=8) or *Il34*^-/-^ rats (n=11). Control rats were given regular water (n=8). *Right panel*. A clinical score of colitis severity was established and followed every day. **(E)** At day 7, rats were sacrificed and the colon length measured. Results are represented as mean ± SEM. Mann Whitney *U* test, a Two-way ANOVA with a Bonferroni posttest for clinical score and Log-rank test for survival analysis, * *p*<0.05, **** *p*<0.0001.

The phenotype of FOXP3+ or FOXP3^-^ CD4+ or CD8+ T cells did not reveal differences in *Il34*^-/-^ rats vs *Il34*^+/+^ (**Supplemental Figures 3D-E**) and we observed similar *in vitro* proliferation capacity for both CD4+ and CD8+ Tregs (**Supplemental Figure 3F**).

Gaining insights toward the role of IL-34 in T cell development/function, we next assessed the impact of the global deficiency in an inflammatory environment using the DSS colitis model. For 7 days, 5.5% DSS given in drinking water to *Il34*^+/+^ and *Il34*^-/-^ rats resulted in an increased severity at all time points in *Il34*^-/-^ animals compared to *Il34*^+/+^ animals (**Figure 2D**). Moreover, analysis of colon length at day 7 showed that the colon of *Il34*^-/-^ rats was shorter than *Il34*^+/+^ and control animals (**Figure 2E**). Altogether, our data demonstrate that the lack of IL-34 upon DSS colitis increases its severity.

### IL-34 deficiency leads to defective suppressive function of CD4+ Tregs *in vivo*

Since we previously showed that around 50% of FOXP3+ CD4+ and CD8+ Tregs express IL-34 (6), we then tested CD4+CD25+ (CD4+ Tregs) and CD8+CD45RC^low/-^ (CD8+ Tregs) Tregs from *Il34*^-/-^ rats for their capacity to control *in vivo* wasting disease by i.v transfer of T CD4+CD45RC^high^ effector cells in *Il2rg*^-/-^ rats (**Figure 3A**) (17). Strikingly, *Il34*^-/-^ CD4+ Tregs were unable to control the development of wasting disease in *Il2rg*^-/-^ animals and death occurred in a similar kinetic than without CD4+ Tregs, in contrast to CD4+ Tregs from *Il34*^+/+^ rats that efficiently controlled wasting disease development (**Figure 3B**), demonstrating the critical role of IL-34 in CD4+ Tregs mediated suppressive activity. Histological analysis of the liver and colon from all groups confirmed the cell infiltration and tissue lesions in *Il2rg*^-/-^ animals injected with CD4+ Teff cells without or with CD4+ Tregs from *Il34*^-/-^ animals, in contrast to animals injected with CD4+ Treg from *Il34*^+/+^ animals (**Figure 3C**). Although we did not observe a protective role of CD8+ Tregs in this model for both *Il34*^+/+^ and *Il34*^-/-^ CD8+ Tregs (**Figure 3D-E**), we observed decreased infiltration in liver and colon in animals injected with CD8+ Tregs from *Il34*^+/+^ animals compared to animals injected with CD4+ Teff cells with or without CD8+ Tregs from *Il34*^-/-^ animals (**Figure 3E**), suggesting that *Il34*^-/-^ CD8+ Tregs were less efficient. To understand how IL-34 deficiency had modified the function of both CD4+ and CD8+ Tregs in *Il34*^-/-^ vs. *Il34*^+/+^ rats, we analyzed their transcriptome using DGE-RNAseq. Intriguingly, the transcriptomic landscape of CD4+ and CD8+ Tregs in *Il34*^-/-^ rats only significantly differed for one gene, *Dnaja1*, a co-chaperone of heat shock proteins (18), identically increased in CD4+ and CD8+ Tregs from *Il34*^-/-^ rats vs CD4+ and CD8+ Tregs from *Il34*^+/+^ rats (**Figure 3F**). Thus, although the role of the increased mRNA expression of *Dnaja1* needs to be explored in future work, the major defect in suppressive capacity observed for Tregs from *Il34*^-/-^ rats is likely not due to a major effect on Tregs direct functions but rather on downstream effects of IL-34 produced by Tregs, mainly through induction of tolerogenic macrophages (4, 6, 9).

**Figure 3.**
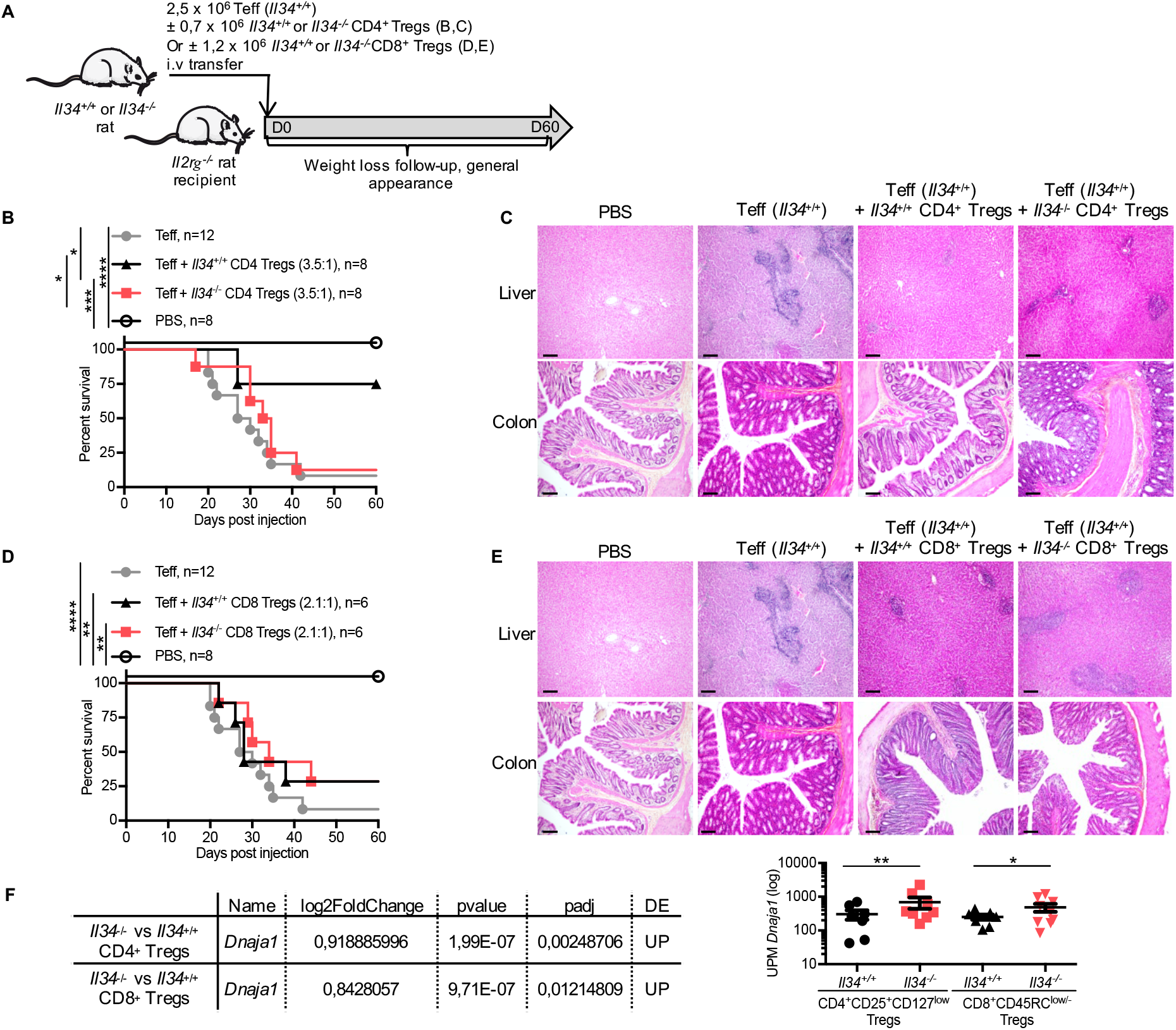
*Il34*^-/-^ CD4 Treg fails to protect from the wasting disease. **(A)** Schematic showing the wasting disease model induction. **(B)** 6 weeks old *Il2rg*^-/-^ rats were injected i.v with 2.5.10^6^ TCRαβ+CD4+CD45RC^high^ Teffs from *Il34*^+/+^ rats (Teffs; n=12) in association or not with TCRαβ+CD4+CD25+CD127^low^ Tregs (CD4+ Tregs; 3.5:1 Teffs:Tregs ratio) from *Il34*^+/+^ (n=8) or *Il34*^-/-^ (n=8) rats in 7 independent experiments. **(C)** Liver and colon sections of injected *Il2rg*^-/-^ rats were stained with HES (original magnification x10, scale bar 20 μm). Data are representative of 6 animals in each group. **(D)** 6 weeks old *Il2rg*^-/-^ rats were injected i.v with TCRαβ+CD4+CD45RC^high^ Teffs from *Il34*^+/+^ rats (Teffs) in association or not with TCRαβ +CD4-CD45RC^low/-^ Tregs (CD8+ Tregs; 2.1:1 Teffs:Tregs ratio) from *Il34*^+/+^ (n=6) or *Il34-*^*/-*^ (n=6) rats in 5 independent experiments. **(E)** Liver and colon sections of injected *Il2rg*^-/-^rats were stained with HES (Original magnification x10). Data are representative of 6 animals in each group. **(F)** 3′ DGE-RNA sequencing analysis was performed on sorted and non-stimulated TCRαβ+CD4+CD25+CD127^low^ Tregs (CD4+ Tregs) and TCRαβ+CD4^-^CD45RC^low/-^ Tregs (CD8+ Tregs) from 8-10 weeks old *Il34*^+/+^ (n=7) and *Il34*^-/-^ rats (n=8-9). The table recapitulates the results of *Dnaja1*, the only gene differentially expressed between *Il34*^-/-^ vs *Il34*^+/+^ CD4+ Tregs and *Il34*^-/-^ vs *Il34*^+/+^ CD8+ Tregs. Mann Whitney *U* test, a Two-way ANOVA with a Bonferroni posttest for clinical score and Log-rank test for survival analysis, * *p*<0.05, ** *p*<0.01, *** *p*<0.001, **** *p*<0.0001.

### IL-34 is expressed by mouse Tregs and *Il34*^-/-^ mice are more prone to autoimmunity

To take advantage of the *Il34*^-/-^ mouse model (12), we first analyzed IL-34 expression by Tregs in *Il34*^+/+^ mice (**Figure 4A**) and observed that CD4+FOXP3+ and CD8+CD45RC^-^ Tregs express significantly more IL-34 than CD4+ and CD8+CD45RC^high^ Teff cells (**Figure 4B**), but at lower levels compared to rats and humans (6). We then assessed susceptibility to autoimmune diseases in *Il34*^-/-^ mice in models of TNBS induced-colitis and experimental autoimmune encephalomyelitis (EAE) triggered by MOG_p35–55_(12). In the TNBS induced-colitis model, we observed a shorter colon length at day 3 in *Il34*^-/-^ mice compared to *Il34*^+/+^ and control mice (**Figure 4C**), indicating a more severe inflammation, however we did not observe significant differences in weight loss (**Supplemental Figure 4A**). We next investigated the impact of IL-34 deficiency in the EAE model since IL-34 could have a protective role through Treg maintenance and microglia development (19, 12, 6). *Il34*^-/-^ mice lost more weight than the controls, but the overall score was not significantly different (**Supplemental Figure 4B**), indicating a mild impact of IL-34 deficiency. On the opposite, when we assessed the therapeutic potential of IL-34 therapy in autoimmunity, we observed that the injection of an adenovirus coding for murine IL-34 in association with a sub-optimal dose of rapamycin led to a delayed EAE development compared to the control groups i.e., null adenovirus associated or not with rapamycin (**Figure 4D**).

**Figure 4.**
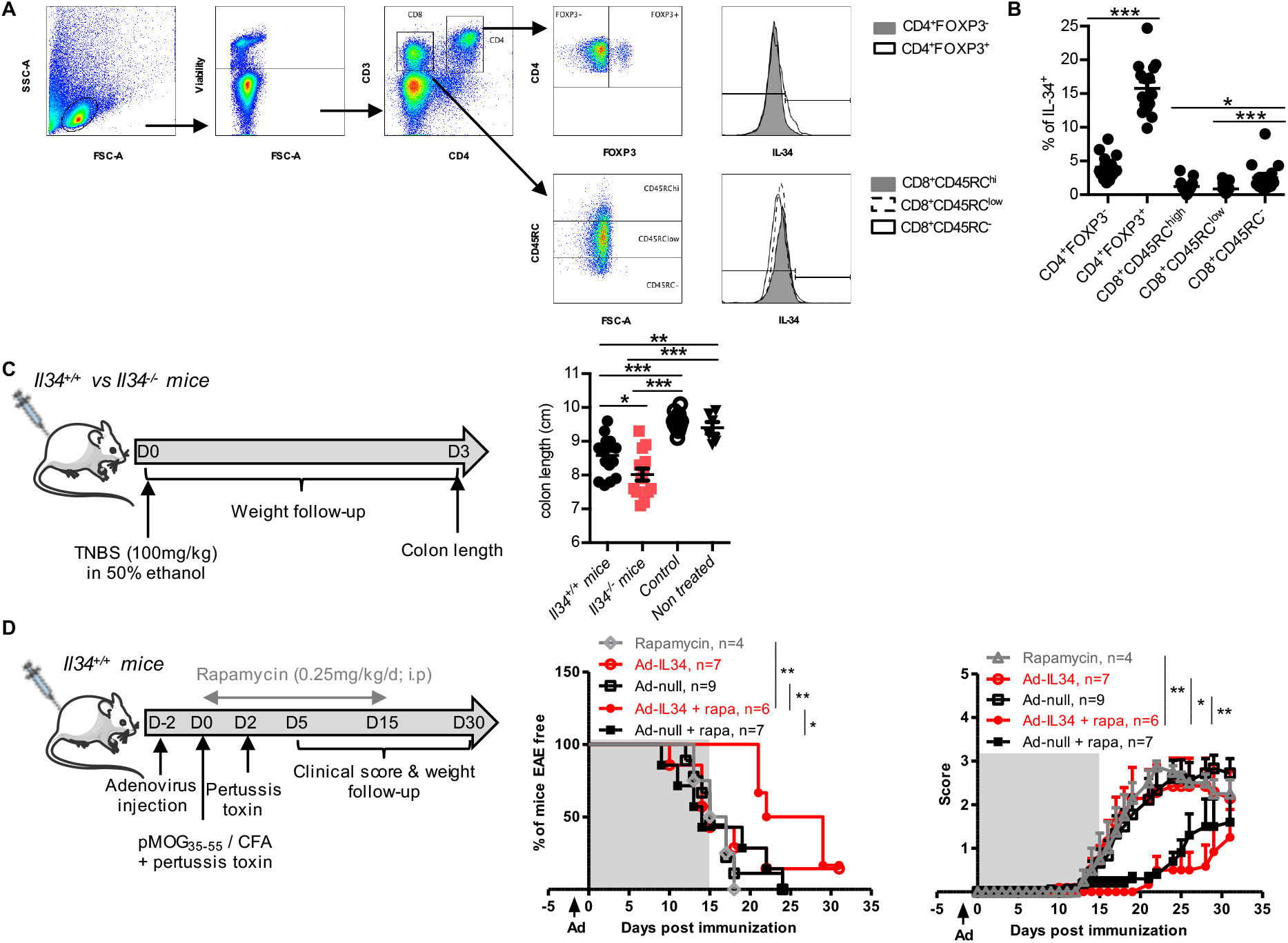
*Il34*^-/-^ mice are more susceptible to autoimmune diseases. **(A)** Gating strategy of IL-34 expression by T cell subsets in mice. **(B)** IL-34 expression was analyzed in CD4+ Teffs (CD4+FOXP3^-^), CD4+ Tregs (CD4+FOXP3+) and CD8+ T cells among the different subsets of CD45RC isoform expression (CD45RC^high/low/-^) (n=15). Negative cells were determined using *Il34*^-/-^ mice. Each dot represents an animal. Results are represented as mean ± SEM. **(C)** Schematic showing the TNBS-induced colitis. The model was performed on *Il34*^+/+^ vs. *Il34*^-/-^ mice (n=11-13) via the intrarectal administration of TNBS (100 mg/kg) in 50% ethanol. Control *Il34*^+/+^ mice were injected with 50% ethanol alone. Mice colon length was measured at day 3 post-injection. Each dot represents an animal. Results are represented as mean ± SEM. **(D)** Schematic showing the IL-34 overexpression in the MOG-induced EAE model. Adult *Il34*^+/+^ mice of 8 weeks-old were i.v injected with an adenovirus coding for mouse IL-34 (Ad-IL-34) or a control (Ad-null) two days before inducing the EAE. Rapamycin was administered or not every day for 15 days at a dose of 0.25mg/kg/d (i.p injection). The development of the disease was followed by a daily weight and scoring assessment. Results are represented as mean ± SEM. Mann Whitney *U* test or two-way ANOVA and a Bonferroni posttest for the score analyses, * *p*<0.05; ** *p*<0.01, *** *p*<0.001, **** *p*<0.0001.

Thus, altogether, IL-34 deficiency in mice leads to an unstable immune phenotype, with increase susceptible to autoimmunity and inflammation in a specific environment. Moreover, supplementing IL-34 is beneficial in neurodegenerative autoimmune diseases in mice.

### hIL-34 protein infusion with a suboptimal dose of rapamycin demonstrates regulatory therapeutic properties

Our data so far present a critical role for IL-34 in Treg-mediated suppression and although we previously showed that IL-34 played a role in a transplantation model in rats and mice (9, 6) the question of whether IL-34 has the capacity to inhibit transplant rejection and induce transplant tolerance in an immune response by human cells still remains. Thus, we used transplantation models in immune humanized immunodeficient NSG mice (**Figure 5A-B**) in which we administered recombinant human IL-34 (rhIL-34) protein using mini-osmotic pumps delivering a constant rate of protein and assessed xenogeneic GVHD or human skin transplant rejection. Administration for 14 days of rhIL-34 protein in association with a sub-optimal dose of rapamycin for 10 days resulted in significant delayed GVHD development with induction of long-term survival in 25% of the recipients (**Figure 5A**) and a significant induction of long-term skin graft tolerance in 50% of the recipients (**Figure 5B**), demonstrating the potential of this therapeutic strategy. Analysis of the proportion of hCD45+ cells showed a decrease in animals treated with rapamycin alone vs. control animals and equal inhibition in animals co-treated with rapamycin and rhIL-34 in the acute GHVD model (**Supplemental Figure 5A**) whereas in the human skin transplantation model the proportion of hCD45+ cells was not modified (**Supplemental Figure 5B**). These results can be explained by the inhibition of T effector development and/or function rather a direct inhibition of human lymphocyte proliferation by IL-34. Whether this is due to an increase in Treg function needs further studies.

**Figure 5.**
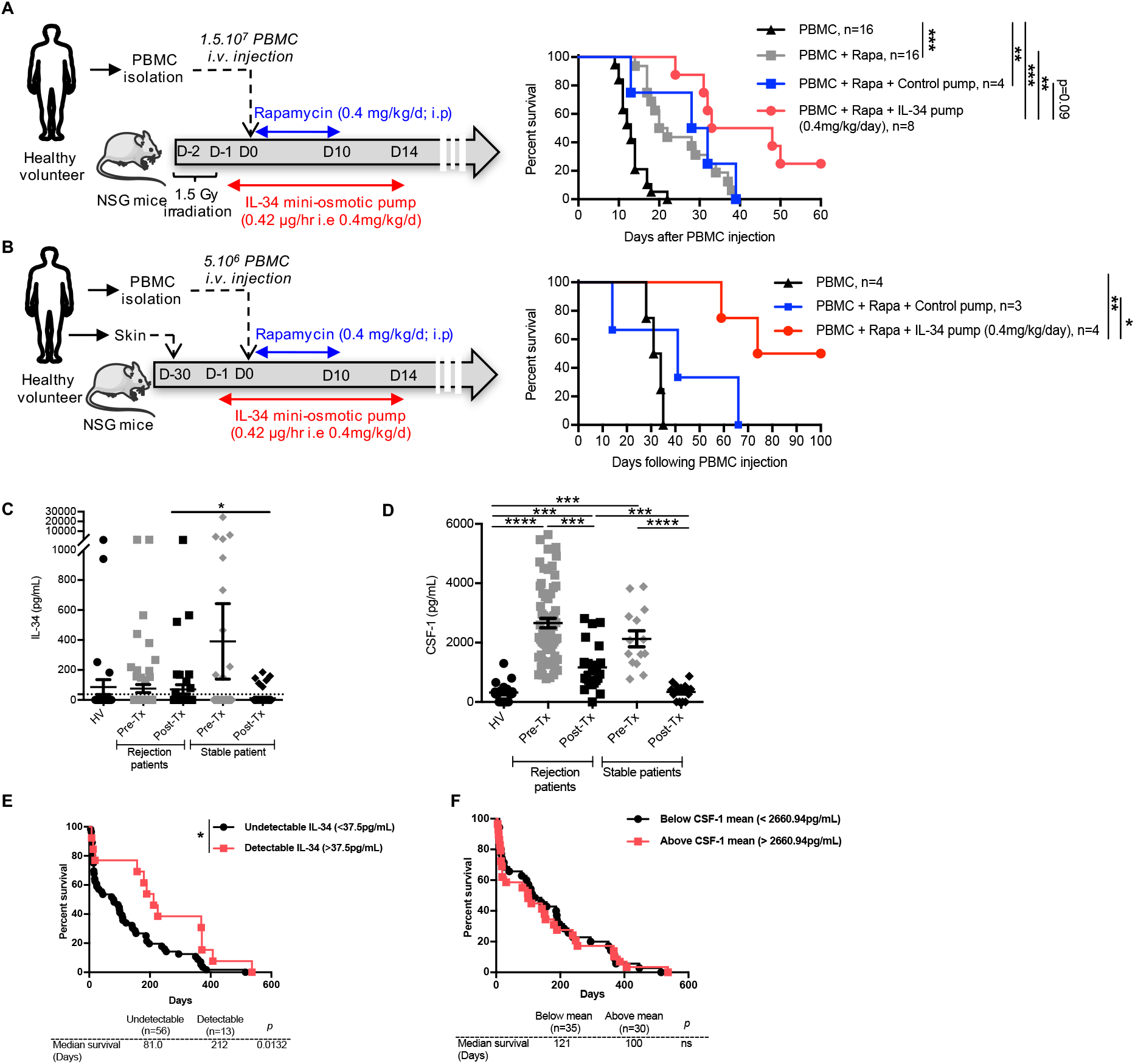
Human IL-34 recombinant protein administration delays GVHD as well as skin allograft rejection in humanized NSG mice and high IL-34 levels predict a better outcome of human kidney transplant. The therapeutic potential of IL-34 was tested in **(A)** GVHD and **(B)** human skin graft allogeneic rejection models in humanized immunodeficient NSG mice. **(A)** IL-34 was administered with a mini-osmotic pump delivering a constant rate for 14 days (0.42 µg/hr) with or without rapamycin (0.4 mg/kg/d for 10 days). **(A)** For GVHD, 1.5.10^7^ fresh PBMC were i.v injected in 1.5 Gy irradiated 8–12-week-old NOD/SCID/*IL2rg*^*™/™*^ (NSG) mice and survival of mice was measured by weight loss. **(B)** For skin allograft rejection, 5.10^6^ fresh PBMC were i.v injected in mice grafted with human skin 4 weeks before. Graft survival was scored on macroscopic signs of rejection from 0 to 5 and considered rejected at a score of 3. **(C)** IL-34 and **(D)** CSF-1 were quantified in the serum of healthy volunteers (n=30 and n=20 respectively) and in patients before (Pre-Tx) and after (Post-Tx) transplantation with a stable graft function or having had ≥ 1 episode of acute rejection within the 18 months following Tx (see Supplemental table 2 for clinical characteristics; Pre-Tx: rejection patients n=71 and n=65, stable patients n=101 and n=14; Post-Tx: rejection patients n=42 and n=22, stable patients n=71 and n=15 for IL-34 and CSF-1 serum levels respectively). **(E-F)** Graft survival of patients with an acute rejection episode occurrence according to pre-Tx IL-34 detectability (undetectable: ≤37.5 vs detectable: ≥37.5 pg/mL) **(E)** or CSF-1 mean expression (below vs above 2660.94 pg/mL) **(F)**. Mann Whitney *U* test and log-rank tests for survival analysis. * *p*>0.05, ** *p*<0.01, *** *p*<0.001 and **** *p*<0.0001.

### Pre-transplantation human IL-34 serum levels, but not of CSF-1, is a prognostic marker of a higher rejection-free episode

We finally assessed the potential use of IL-34 as abiomarker in organ transplantation compared to CSF-1 using serum samples extracted from the DIVAT cohort biocollection (**Supplemental table 2**). We quantified IL-34 and CSF-1 levels by ELISA in sera from a cohort of kidney transplant patients before (Pre-Tx) and within 18 months after kidney transplantation (Post-Tx). Serum levels were also compared according to graft outcome within the observation period of 18 months, i.e. either patients with a stable graft function, free of rejection episodes or with episodes of acute rejection, to identify stable patients vs rejection patients (**Figure 5C-F**). We first determined the detectability of IL-34 based on the detection threshold: 37.5 pg/mL i.e. the lowest concentration of the standard curve provided by the manufacturer in the sera of patients. IL-34 serum levels were detectable (≥ 37.5 pg/mL) in around 22% of patients in the different groups (**Supplemental Figure 6A**) with increased levels in patients who had experienced a rejection compared to stable patients at the post-transplantation period (**Figure 5C)**. In contrast, CSF-1 was detectable in most individuals and levels were higher in pre-transplantation compared to post-transplantation as well as in healthy volunteers. CSF-1 levels were also higher in post-transplantation with rejection vs. stable patients (**Figure 5D**). We further analyzed the correlation between IL-34/CSF-1 and the clinical outcome to determine the possibility of using IL-34 or CSF-1 as a prognosis biomarker. We split patients with rejection episodes in two groups based on IL-34 detectability (≤37.5 pg/mL vs ≥ 37.5 pg/mL) (**Figure 5E and Supplemental Figure 6A**) or CSF-1 mean expression for all patients (≤ 2660.94 pg/mL vs ≥ 2660.94 pg/mL) (**Figure 5F**) in serum before transplantation and analyzed the graft survival period before acute rejection episode occurrence. Interestingly and unlike CSF-1, the proportion of patients without rejection episodes in patients with IL-34 levels ≥ 37.5 pg/mL before transplantation was significantly higher than the one of patients with IL-34 below this level. This statistical correlation was specific when measured before transplantation since IL-34 levels in serum after transplantation did not predict graft outcome (**Supplemental Figure 6B-C**). Our results thus suggest that IL-34 can act as a prognosis biomarker before transplantation.

## Discussion

Here, we demonstrated IL-34 as a critical actor of Treg-mediated suppression, playing an important role in homeostasis and control of immune responses. IL-34 expression in the T cell lineage seems to be restricted to FOXP3+ CD4+ and CD8+ Tregs (6, 9) acting as a significant suppressor component of their activity. IL-34 is also expressed by other cell subsets such as neurons playing as well a protective role for microglia (20, 21).

At steady state, the overall phenotype of *Il34*^-/-^ rats remained similar to the one of *Il34*^+/+^ mice (12, 19) with no evident developmental or pathological signs with the exception of a decrease in microglia. There has not been a thorough description of the distribution and function of other immune cell subsets in *Il34*^-/-^ mice or on the response to inflammatory/autoimmune stimuli, as described here for *Il34*^-/-^ animals. The increased expression of auto-antibodies in *Il34*^-/-^ rats is intriguing suggesting a role for IL-34 the regulation of antibody production. These autoantibodies include anti-IFN-I antibodies that in addition to the role of proinflammatory monocytes/macrophages in the cytokine storm leading to severe acute respiratory distress syndrome and multi-organ pathology observed in SARS-CoV-2 infection could suggest a role for IL-34 in COVID-19 patients (22, 23).

IL-34 deficiency in rats led to a mild immune phenotype *in vivo* at resting state, probably due to CSF-1 over-expression that may partially compensate IL-34 deficiency in their common overlapping functions, particularly in monocyte and macrophage differentiation. These results suggest a negative regulation of CSF-1 by IL-34, complementary to the positive regulation of CSF-1 on IL-34 previously described (24). Despite the fact that CSF-1 could replace activities of IL-34, we still observed some features of a disturbed immune system and differences on the CD8+ T cells lineage distribution pattern. However, upon challenges, we revealed an increased susceptibility to colitis. Even though we did not find any perturbation in myeloid distribution in the gut at steady state (data not shown), one explanation could be that the inflammatory environment could not be counterbalanced by the action of IL-34 on macrophage polarization leading to a greater severity in deficient animals. In the EAE model, the action of IL-34 can be mediated either through the development of microglia (5) or through Treg activity both in the CNS and periphery. As we did not find differences in EAE development when comparing *Il34*^*+/-*^ to *Il34*^-/-^ mice, this would suggest a mild role of IL-34. However, overexpression of IL-34 was beneficial and significantly delayed EAE development showing a therapeutic potential of IL-34 in the initial phases of inflammatory CNS diseases. When we specifically studied the impact of the deficiency in the Treg subsets, we revealed a clear defect in the IL-34 deficient CD4+ Tregs suppressive function where they fail to protect from the wasting disease. This could be due to an insufficient differentiation of M2-like macrophages. This is reminiscent of our observations that IL-34 induces immune tolerance in a model of organ transplantation through M2 macrophages and downstream induction of Treg (9), creating a positive regulatory loop. Interestingly, transcriptomic analysis of T cells revealed the upregulation only of *Dnaja1* in both *Il34*^-/-^ CD4+ and CD8+ Tregs compared to *Il34*^+/+^ rats. *Dnaja1* encodes for a protein belonging to the heat shock protein 40 (HSP40) acting as a HSP70 co-chaperone. One report has demonstrated a regulatory role of Dnaja1 where it inhibits T cell proliferation and induces IL-10 production by PBMC in rheumatoid arthritis patients (25). Further work needs to be done in order to comprehend the link between IL-34 and Dnaja1.

IL-34 upregulation has been associated to certain but not all auto-immune diseases. However, the exact contribution of IL-34 in these diseases has not been elucidated since IL-34 upregulation could be a compensatory tolerogenic mechanism, thus a consequence rather than a cause (1, 2). Moreover, it is important to take into consideration that although IL-34 and CSF-1 share the CSF-1R that could explain results through M2 cells, IL-34 has two other receptors or co-receptors, PTPζ and CD138, expressed on other cell types, that could also participate on its final effects (2, 5). IL-34 was shown to delay solid organ graft rejection in animal models and associated with a delayed liver rejection in human (6, 11). The delay in skin rejection and GVHD in the NSG immune humanized models of the present manuscript are the first ones showing a potent suppressive effect on human immune responses *in vivo*. As monocytes in NSG mice are only detected in the first few days post-hPBMC injection and rapamycin is needed in this context and in the allotransplantation model in rat (6), we believe, in accordance with previous results (9) that IL-34 first acts on monocytes to differentiate them in M2-like macrophages which will allow to expand Tregs, creating a regulatory environment. Along this line of inhibition of allogeneic immune responses by IL-34, detection of IL-34 in the sera of patients before kidney transplantation defined patients that have a higher probability of having a longer rejection-free post-transplantation period. Thus, and unlike CSF-1, detectable IL-34 levels in the serum before transplantation could serve as a biomarker of better kidney transplant outcome. Still, this finding needs to be confirmed in a larger cohort and confronted to other cohorts of transplanted patients.

Overall, we show that IL-34 is crucial to CD4+ Tregs suppressive function as well as its deficiency led to increased susceptibility in some autoimmune processes. In human, administration of IL-34 represents a new therapeutic strategy for treatment of autoimmune diseases, GVHD and transplant rejection.

## Methods

### DIVAT cohort of kidney transplanted patients

Serum samples from kidney +/-pancreas transplanted patients were obtained thanks to the “Données Informatisées et VAlidées en Transplantation” DIVAT Biocollection (www.divat.fr, French Research Ministry: RC12_0452, last agreement No 13 334, No CNIL for the cohort: 891735) collected between 2004 and 2012, aliquoted and stored at -80°C. All recruited patients gave signed informed consent. Supplemental Table 2 recapitulates the clinical characteristics of the selected cohort.

### Animals

Sprague-Dawley (CD) rats were purchased from Charles River France (L’Arbresle, France) and C57Bl/6 mice from Janvier Labs (Le Genest-Saint-Isle, France). The 8–12-week-old NOD/SCID/*Il2rg*^*™/™*^ (NSG) mice were bred in our own animal facilities in SPF conditions. *Il34*^LacZ/LacZ^ mice were kindly provided by Marco Colonna (Washington University, St. Louis, MO) (12). *Il2rg*^-/-^ rats were kindly provided by TRIP Platform (Nantes, France) and have been previously described (26). The animals were housed in a controlled environment (temperature 21 ± 1 °C, 12-hour light/dark cycle).

### Generation and genotyping of Il34^-/-^ SPD rats

rIL34-sgRNA4 (TGTACTGCAGCTTGCCCCGA) was designed and generated as previously described (27). Prepubescent females (4–5 weeks old) were injected with 25 IU pregnant mare serum gonadotropin (Intervet) and followed 48 hours later with 30 IU human chorionic gonadotropin (Centravet, Plancoet, France) before breeding (27). Fertilized one-cell stage embryos were collected for subsequent microinjection using a previously published procedure (27, 28). sgRNA (10ng/µl) and Cas9 mRNA (50ng/µl) (27) were co-microinjected into the cytoplasm and nucleus of one-cell-stage fertilized embryos. Surviving embryos were implanted in the oviduct of pseudo-pregnant females (0.5dpc) and allowed to develop until birth. For genotyping of animals, ear biopsy specimens from 8-to 10-d-old rat pups were digested in 250 µl of tissue digestion buffer (Tris-HCl 0.1 mol/l [pH 8.3], EDTA 5 mmol/l, SDS 0.2%, NaCl 0.2 mol/l, proteinase K 100 µg/ml) at 56°C overnight. PCR amplification was performed on diluted lysis product (1:20 dilution) and 25 µl of PCR reaction mix according to the manufacturer instruction (Herculase II Fusion DNA Polymerase, Agilent Technologies, Les Ulis, France) using the following PCR primers: forward 5’-AGGTGGAGTACAGACACAGT-3’ and reverse 5’-AGATAAGAGGTGGGAGTGAGC-3’. The following amplification program was used: 1 cycle at 95°C for 5 min, 1 cycle at 62°C for 2min, 35 cycles at 72°C for 30 s, 95°C for 10 s, and 60°C for 10 s, followed by 1 cycle at 72°C for 3 min using a Veriti Thermal Cycler (Applied Biosystems, Foster City, CA, USA). The PCR products were analyzed by heteroduplex mobility assay using microfluidic capillary electrophoresis system caliper LabChip GX (PerkinElmer, Villebon-sur-Yvette, France). The homozygous *Il34*^-/-^ specimen were identified from *Il34*^+/+^ littermate as previously described (13).

### Bone density

Femurs of *Il34*^+/+^ and *Il34*^-/-^ rats were fixed 24h in PFA 4%, then trabercular bone mineral density (BMD) and cortical tissue mineral density (TMD) were analyzed using the high-resolution SkyScan-1076 X-ray microCT system (SkyScan, Kartuizersweg, Belgium).

### ELISA

Rat CSF-1 was quantified using a rat M-CSF ELISA (MyBioSource, San Diego, CA, USA). Human CSF-1 and IL-34 human serum levels were quantified using ELISA kits (both R&D systems, Mountain View, CA, USA), according to manufacturer’s instructions. Anti-dsDNA antibodies were quantified by coating 100 µg/mL of salmon sperm DNA (Invitrogen) in a 96-well flat-bottom plate (Thermo Fisher Scientific) in PBS overnight at 4 °C. After 5 washes in PBS-Tween 20 0.05%, saturation with PBS-BSA 5%-FCS 2% was performed for 1h30 at 37 °C. Sera from *Il34*^+/+^ or *Il34*^-/-^ rats were serially diluted in PBS-BSA 1% (1/2, 1/20, 1/200 and 1/2000) and incubated for 1 h at 37°C. After 10 washes with PBS-Tween 20 0.05%, an HRP-conjugated goat anti-rat IgG (heavy and light) (Jackson Immunoresearch) was used as secondary Ab (1 h, 37 °C). 10 washes in PBS-Tween 20 0.05% and the reaction was visualized by the addition of chromogenic substrate (TMB, BD Biosciences). A stop solution (H_2_SO_4_) was added and absorbance at 450 nm was measured with reduction at 630 nm using an ELISA plate reader. Results are expressed in optical density (O.D). IgM, IgG, IgA and IgE were measured in the serum as previously described (26).

### Luminex

Rat serum concentrations of TGFβ1, TGFβ2, TGFβ3, Eotaxin, MIP-2, MIP-1α, IL-13, IL-1β, IL-1α, IL-2, IL-4, IL-5, IL-6, IL-10, IP-10, IL-12p70, IL-17A, G-CSF, GM-CSF, TNFα, IFNγ, GROα, MCP-1, MCP-3 and RANTES were quantified using a multiplex kit (MILLIPLEX MAP; Merck, Burlington, MA and ProtocartaPlex; ThermoFischer, Waltham, MA) according to manufacturer’s instructions.

### Luciferase

*Immunoprecipitation Systems (LIPS)*. LIPS was used for the detection of autoantibodies against cytokines as previously described (29, 30).

### qPCR

Total RNA was isolated from cells using TRIzol reagent (Invitrogen) or a RNeasy Mini Kit (QIAGEN, Hilden, Germany). Total RNA was isolated from organs (spleen, liver and brain) by crushing with Ultra-Turrax (IKA, Staufen, Germany) using TRIzol reagent (Invitrogen, Carlsbad, CA) and RNA was reverse transcribed with random primers and M-MLV reverse transcriptase (Life Technologies, Carlsbad, CA) according to manufacturer’s instructions. For rat *Il34* qPCR, TaqMan (Thermofischer, Waltham, MA) probes were used (Rn01432380_m1; Thermofischer, Waltham, MA). The reaction was performed on the Applied Biosystems StepOne system (Life Technologies, Carlsbad, CA). Thermal conditions were as follows: 3 seconds at 95°C, 30 seconds at 60°C, and 15 seconds at –5°C of the melting temperature with a final melting curve stage. Calculations were made by the ddCt method.

### Histology and Immunofluorescence

Rat brains were fixed for 24h using PFA 4% and conserved by increasing sucrose concentrations and frozen. Sections were performed from paraffin-embedded tissues and frozen brain tissues. For histology analysis, slides were stained with Hematoxylin Eosin Saffron (HES) and analyzed by an automated tissue slide scanner (Hamamatsu NanoZoomer Digital Pathology system, Japan) and by confocal microscopy (Nikon A1 RSi, Tokyo, Japan). For brain immunofluorescence, slides were incubated with a purified anti-CD11b/c (a list of the clones and suppliers of all mAbs used in the study in Supplemental Table 1) and the staining was revealed using a goat anti-mouse IgG (H+L)-AF488 (Invitrogen, Carlsbad, CA). After staining, slides were mounted with Prolong Gold Antifade Reagent with DAPI (Invitrogen, Carlsbad, CA) before analysis with confocal microscopy. The percentage of positive staining was analyzed using ImageJ software.

### Cell isolation

Rat spleen was digested by collagenase D (Roche) for 30 min at 37°C; the reaction was stopped by adding 0.01 mM EDTA. Mice spleen and rats thymic cells and lymph nodes were isolated by crushing with PBS. Red blood cells were lysed using a lysis solution (8.29 g NH4Cl, 1 g KHCO3, 37.2 mg EDTA/1 l deionized water [pH 7.2–7.4]). Rat TCRαβ-SIRPα+, TCRαβ+CD4+CD25^−^, TCRαβ+CD4+CD45RC^high^, TCRαβ+CD4^-^CD45RC^high^, TCRαβ+CD4+CD25+CD127^low^ and TCRαβ+CD4^-^CD45RC^low/-^ cells were sorted using a FACS ARIA II (BD Biosciences, Mountain View, CA). Monoclonal antibodies are listed in Supplemental Table 1.

### Phenotypic analysis

Cellular phenotype was analyzed on spleen, thymus and blood using the antibodies listed in the Supplemental Table 1. Cells were first gated on their morphology, exclusion of singlets and dead cells by staining with fixable viability dye, eF450 or eF506 (Thermofischer, Waltham, MA), then cells were gated on the expression of CD45. Rat subsets were identified as follow: DN (CD4^-^CD8^-^), DN1 (CD4^-^CD8^-^CD25^-^CD44+), DN2 (CD4^-^CD8^-^ CD25+CD44+), DN3 (CD4^-^CD8^-^CD25+CD44^-^), DN4 (CD4^-^CD8^-^CD25^-^CD44^-^), SP CD4+ (CD4+CD8^-^), SP CD8+ (CD4^-^CD8+), IgM and IgD expression were analyzed among CD45R+CD45RA+ cells, pDC (TCRαβ^-^CD4+CD45R+), cDC (TCRαβ^-^CD4^+/-^CD103+), NK (SIRPa^-^TCRαβ^-^CD161^++^), NKT (SIRPa^-^TCRαβ+CD161+), granulocyte (TCRαβ^-^HIS48+RP-1+), macrophage/monocyte (CD68+), M1 (CD68+CD163^-^) and M2 (CD68+CD163+). Mouse subsets were identified as follow: CD4 Teff (CD3+CD4+FOXP3^-^), CD4 Treg (CD3+CD4+FOXP3+), CD8 Teff (CD3+CD4^-^CD45RC^high^), CD8+CD45RC^int^ (CD3+CD4^-^ CD45RC^int^) and CD8+CD45RC^-^ (CD3+CD4^-^CD45RC^-^). For stimulation, splenocytes were incubated with PMA (50 ng/ml) and ionomycin (1 µg/ml) for 4 h in the presence of Brefeldin A (10 µg/ml). Cells were permeabilized with Fix/Perm kit (Thermofischer, Waltham, MA). Abs were used to stain cells, and fluorescence was measured with a BD FACSCanto II flow cytometer (BD Biosciences, Mountain View, CA), and FlowJo software was used to analyze data.

### Proliferation assay

Sorted rat TCRαβ+CD4+CD25^−^ and TCRαβ+CD4^-^CD45RC^high^ Teffs, TCRαβ+CD4+CD25+CD127^low^ and TCRαβ+CD4^-^CD45RC^low/-^ Treg cells from *Il34*^+/+^ and *Il34*^-/-^ rats were plated in duplicate or triplicate with stimulating coated αCD3 (G4.18 at 1, 0.5 and 0.25 µg/mL) and soluble αCD28 (JJ319 at 10 µg/mL) mAbs in 100 µL or 200 µL complete RPMI-1640 medium in round-or V-bottomed 96-well plates, at 37°C and 5% CO_2._ Proliferation of CFSE-labeled TCRαβ+CD4+CD25^−^, TCRαβ+CD4^-^CD45RC^high^, TCRαβ+CD4+CD25+CD127^low^, TCRαβ+CD4^-^CD45RC^low/-^ T cells were analyzed by flow cytometry 2 or 3 days later by gating on TCRαβ+CD4+ cells among living cells (DAPI^-^).

### DSS colitis in rats

DSS (TdB labs, Sweden) was dissolved in drinking water at 5.5% and given to 9-weeks old male rats for 7 days. Weight loss and a disease activity index (DAI=clinical score) were assessed every day. Score is defined as follow: stool consistency: 0 (normal), 2 (loose stool), and 4 (diarrhea); and bleeding: 0 (no blood), 2 (visual pellet bleeding), and 4 (gross bleeding, blood around anus). A day 7, animals were sacrificed and colon length was measured to estimate the inflammation.

### TNBS-induced colitis in mice

The model of colitis was induced by rectal administration of TNBS (100 mg/kg in 0.1 mL; Merck, Burlington, MA) in 50% ethanol. Animals were weighed daily and were sacrificed 3 days post-injection to measure colon length.

### Experimental autoimmune encephalomyelitis (EAE)

Eight-week-old *Il34*^+/+^ or *Il34*^*LacZ*/*LacZ*^ C57BL/6J female mice (12) were immunized by subcutaneous injection of 100 µL of an emulsion containing 200 µg of MOG_p35–55_(GeneCust, Boynes, France) in complete Freunds adjuvant (Merck, Burlington, MA) supplemented with 400 µg heat-killed *M. tuberculosis* (Thermofischer, Waltham, MA). 200ng of *B. pertussis* toxin (Merck, Burlington, MA) in 100 µl of PBS was injected intraperitoneally on the same day and 2 days after immunization. Adenovirus, coding for murine IL-34 or null, were injected i.v at 4.10^9^IP/100uL/mice two days before the immunization. Rapamycin was i.p injected every day for 15 days at 0.25mg/kg (Rapamune, Pfizer). Animals were monitored daily from day 5 and scored on a 5-point scale as follows: 0, no symptoms; 0.5, tip of tail is limp; 1, loss of total tail tonus; 1.5, loss of total tail tonus and hind leg inhibition; 2, loss of total tail tonus and weakness of hind legs; 2.5, loss of total tail tonus and dragging of hind legs; 3, loss of total tail tonus complete and paralysis of hind legs; 3.5, loss of total tail tonus and paralyzed hind legs are together on one side of the body; 4, loss of total tail tonus, hind leg and partial front leg paralysis; 4.5, loss of total tail tonus, hind leg and front leg paralysis, 5: mouse is found dead due to paralysis. Due to ethical considerations mice were sacrificed when they reached grade 4 for 24 h.

### Wasting disease. Il2rg^-/-^

SPD rats aged of 6 weeks were injected via the tail vein with 2.5.10^6^ sorted TCRαβ+CD4+CD45RC^high^ Teff cells from *Il34*^+/+^ rats in association or not with sorted TCRαβ+CD4+CD25+CD127^low^ or TCRαβ+CD4^-^CD45RC^low/-^ Tregs from *Il34*^+/+^ or *Il34*^-/-^ rats at a Teffs:Tregs ratio of 3.5:1 and 2.1:1 respectively. A control with a PBS injection was also performed.

### 3’ Differential Gene Expression (DGE) RNA-sequencing

Total mRNA from sorted TCRαβ+CD4+CD25^−^ and TCRαβ+CD4^-^CD45RC^high^ Teffs, TCRαβ+CD4+CD25+CD127^low^ Tregs, TCRαβ+CD4^-^CD45RC^low/-^ T cells from *Il34*^+/+^ and *Il34*^-/-^ rats were extracted using RNeasy Mini Kit (QIAGEN, Hilden, Germany) and protocol of 3’ DGE RNA sequencing was performed as previously described (16). The differential expression p values were processed with DESeq2 (31).

### Immune humanized mouse models

For xenogeneic graft-versus-host-disease (GVHD) model, 1.5×10^7^ fresh human PBMC were intravenously injected in 1.5 Gy irradiated NSG mice as previously described (10). Human PBMC engraftment was monitored in blood and GVHD development was characterized by ≥20% body weight loss. For the skin rejection model, human skins were obtained from healthy donors from abdominoplasty surgery and transplantation was performed as previously described (10). 5.0×10^6^ PBMC allogeneic to the graft were i.v injected. A graft rejection score was established from 0 to 5 based on macroscopic observations: 1: the skin starts to peel off; 2: thick skin; 3: scab; 4: edges start to take off; 5: the skin is entirely gone. Osmotic pumps (Alzet, model 1004, Cupertino, CA) were filled with rhIL-34 (0.42µg/h, i.e. 0.4 m/kg/d; Thermofisher, Waltham, MA or Preprotech, Neuilly-Sur-Seine, France) and placed i.p on the day before the injection of the PBMC. Rapamycin (0.4 mg/kg/d for 10 days, Rapamune, Pfizer) was injected intraperitoneally.

### Study Approvals

All animal care procedures were approved by the Animal Experimentation Ethics Committee of the Pays de la Loire region, France, in accordance with the guidelines from the French National Research Council for the Care and Use of Laboratory Animals (permits numbers CEEA-PdL-n°6, APAFIS #12377, #20640, #2162, #0692, #18724 and #27925). For the DIVAT cohort, all recruited patients gave signed informed consent.

### Statistics

Mann-Whitney *U* test was used for qPCR, FACS, positive area in immunohistofluorescence and ELISA analysis. Mantel Cox Log Rank test was used to analyze survival curves. Two-way ANOVA and a Bonferroni posttest were used to compare weight loss and clinical score between groups. Adapted controls were performed together with the test conditions. Animal numbers were determined with ethical committee agreement.

## Author contributions

A.F. contributed to data collection, experimentation, analysis and writing of the manuscript.

A.S., S.B., L.T., N.V., R.H, H.R., L.F., P.P., S.R. and F.B. contributed to data collection, experimentation and analysis.

C. U., J.-M.H., S.R., S.M., LT and IA generated the IL-34 deficient rats.

F.D, M.G. and M.C. contributed with critical reagents.

I.A. and C.G. led and conceived the project, obtained funding, analyzed the data and wrote the manuscript.

## Acknowledgments

We thank Séverine Battaglia (Inserm UMR1238, Nantes, France) for performing the Micro-CT analysis and Ms Maire Pihlap for the LIPS analysis. We thank the biological resource center for biobanking of Nantes Hotel-Dieu University hospital (CRB Nantes, F-44093, France -BRIF: BB-0033-00040) for their help in the preparation of the samples. We thank Laure-Hélène Ouisse (ITUN CRTI UMR1064, Nantes, France) for the Ig quantification. This work was partially funded by Labex IGO project (n°ANR-11-LABX-0016-01). Labex IGO is part of the « Investissements d’Avenir » French Government program managed by the ANR. This work was funded by the Agence Nationale de la Recherche ANR-17-CE18-0008, the Fondation du rein “Don de Soi – Don de Vie 2017 FdR-Trans-Forme/FRM” and the Agence de la Biomedecine. This work was also realized in the context of the support provided by the Fondation Progreffe.

## Data availability statement

The data that support the findings of this study are available from the corresponding author upon reasonable request.

