## Supplemental_Figures_Tables for "IL-34 deficiency impairs FOXP3^+^ Treg function and increases susceptibility to autoimmunity"

Freuchet et al. Suppl. Figure 1

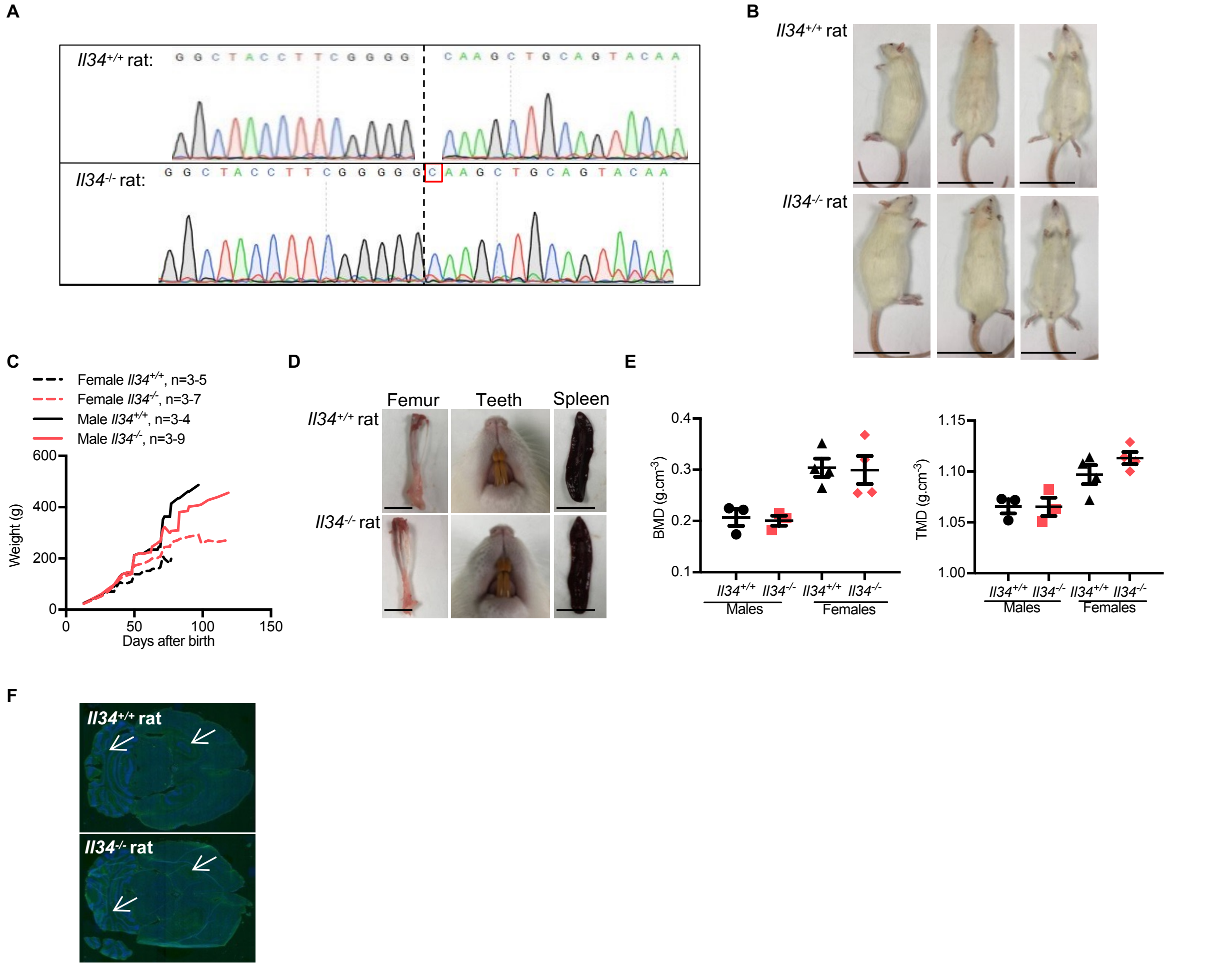


**Supplemental Figure 1. Generation of *Il34*$^{-/-}$ rats and characterization**. **(A)** Amplicons of *Il34*$^{+/+}$ and *Il34*$^{-/-}$ rats were sequenced showing a C insertion in mutated animals, leading to a shift in the open reading frame and a premature STOP codon (not shown in the electrophorogram). **(B)** Photographs represent the general appearance of the rats at 6 months old (scale bar, 10 cm). **(C)** Littermate *Il34*$^{+/+}$ rats (males, n=3-4 and females, n=3-5) and *Il34*$^{-/-}$ rats (males, n=3-9 and females, n=3-7) were weighed for 100 days. Results are expressed as mean ± SEM. **(D)** Representative photographs show the bone aspect, teeth formation and spleen length (Scale bar, 1 cm). **(E)** Femurs of 8 weeks old males (n=3) and females (n=4) *Il34*$^{+/+}$ or *Il34*$^{-/-}$ were fixed in 4% formol and then analyzed by µCT for bone mineral density (BMD) and tissue mineral density (TMD). **(F)** Representative photographs show the morphological aspect of *Il34*$^{+/+}$ (top) and *Il34*$^{-/-}$ (bottom) brains. White arrows indicate the zoom in of the hippocampus and cerebellum showed in Figure 1D. Mann Whitney *U* test or two-way ANOVA and a Bonferroni posttest, ns $p>0.05$.

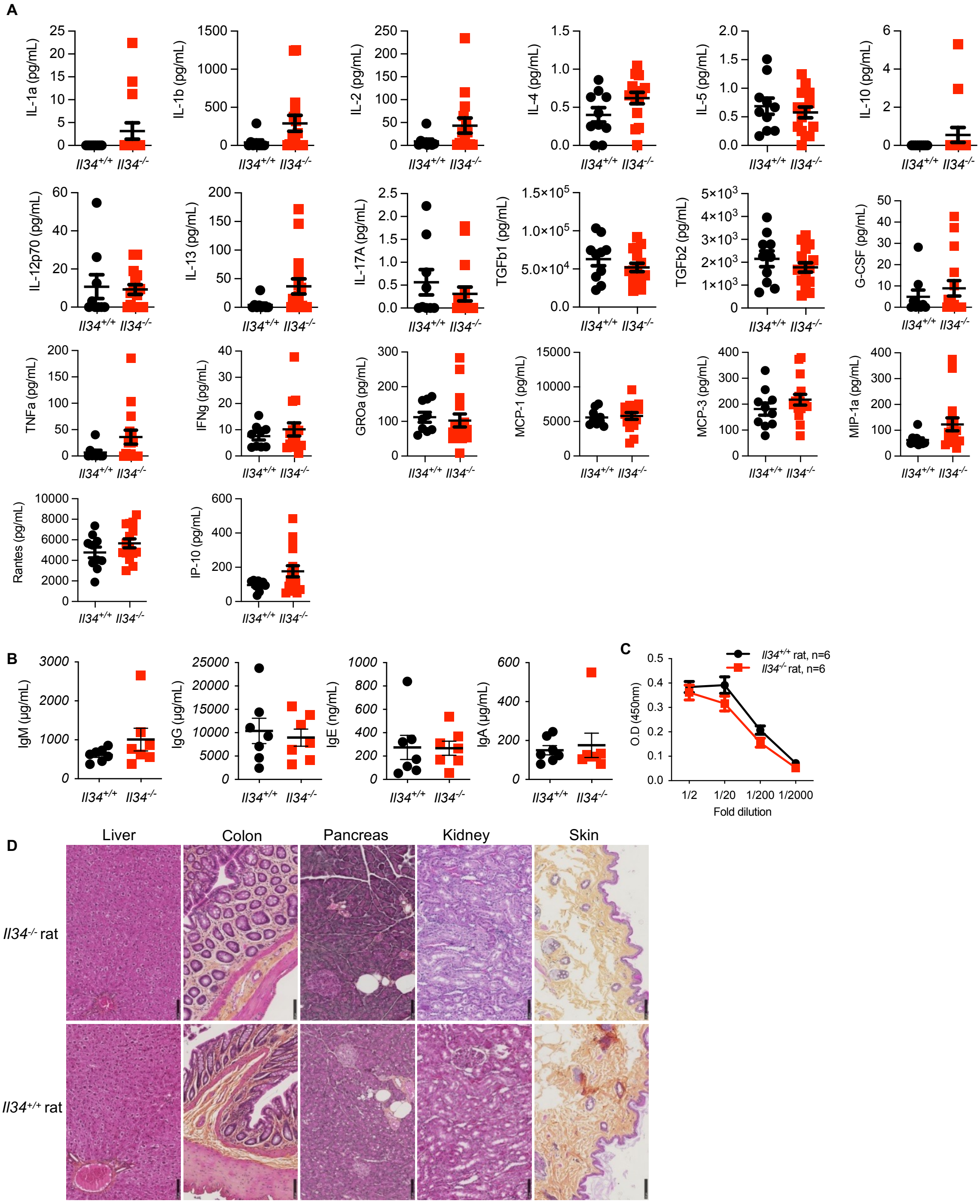


**Supplemental Figure 2. Quantification of cytokines, chemokines, growth factors, antibodies and histological analysis of *Il34*$^{+/+}$ vs *Il34*$^{-/-}$ rats (A)** Plasma levels of TGFβ1/2, IL-1α/β, -2, -4, -5, -6, -10, -12p70, -13, -17A, G-CSF, GM-CSF, TNFα, IFNγ, GROα, MCP-1, -3, MIP-1α, Rantes and IP-10 were quantified in 4 months old *Il34*$^{+/+}$ (n=9) and *Il34*$^{-/-}$ animals (n=14). GM-CSF and IL-6 were not detectable. **(B)** IgM, IgG, IgE and IgA were quantified in the plasma of *Il34*$^{+/+}$ (n=7) and *Il34*$^{-/-}$ rats (n=7). **(C)** Anti-double strain DNA antibodies were measured at different dilutions (1/2, 1/20, 1/200 and 1/2000) by ELISA (n=6 per group). Results are represented as mean ± SEM. **(D)** Liver, colon, pancreas, kidney and skin sections of *Il34*$^{+/+}$ and *Il34*$^{-/-}$ rats at 6 months old were stained with HES. Original magnification × 20, scale bar 100 µm. Mann Whitney *U* test or two-way ANOVA and a Bonferroni posttest, ns $p$>0.05.

Freuchet et al. Suppl. Figure 3

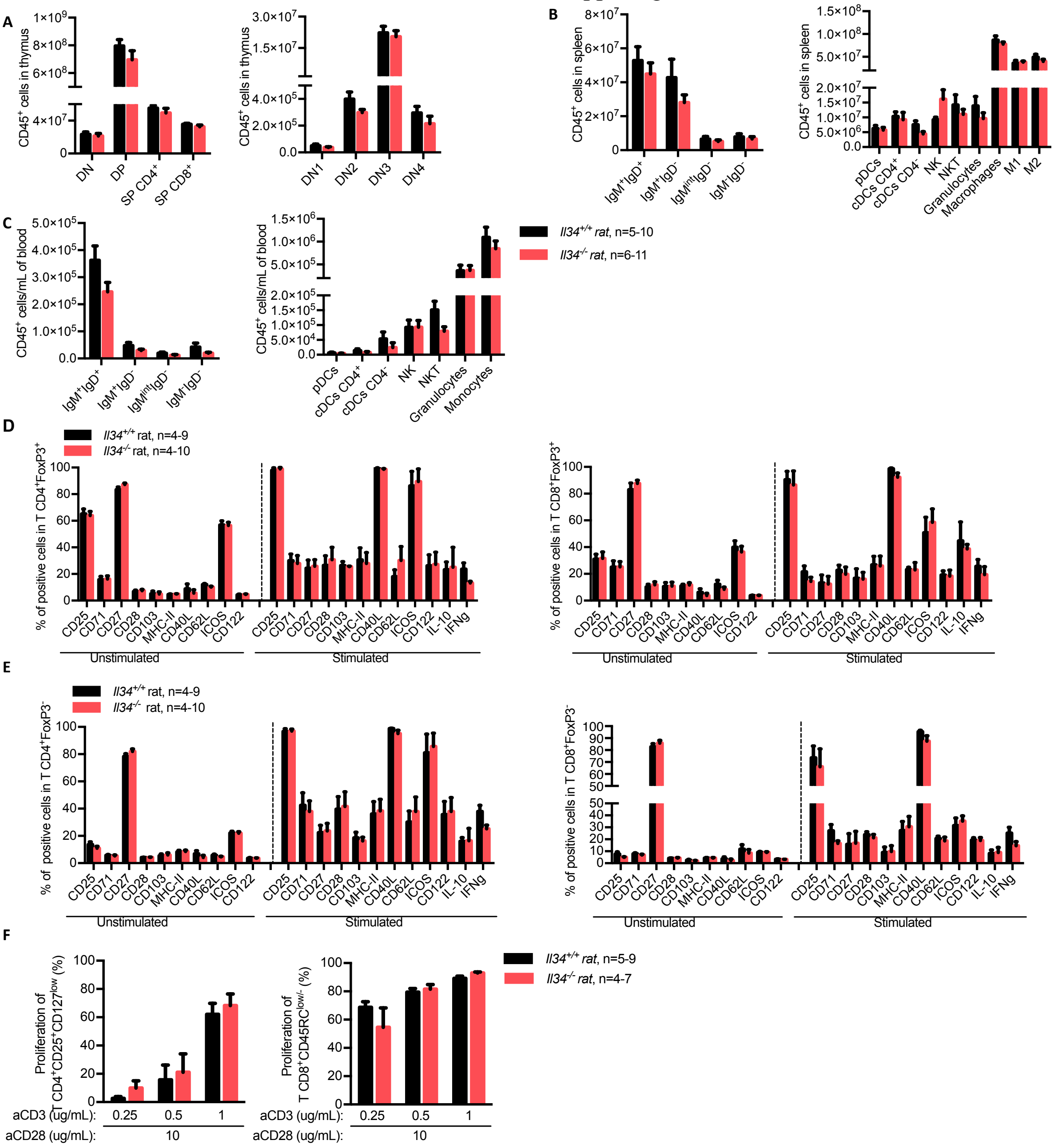


**Supplemental Figure 3. Immune cell populations are not affected by IL-34-deficiency at resting state. (A)** Absolute numbers of thymic cell subsets were analyzed in 8 weeks old *Il34*$^{+/+}$ (n=7) and *Il34*$^{-/-}$ (n=8) rats using markers described in *Supplemental Methods*. Absolute numbers of B, DC, NK, NKT, granulocyte and monocyte/macrophage cell subsets were analyzed in spleen **(B)** and blood **(C)** of >10 months olds *Il34*$^{+/+}$ (n=4-9) and *Il34*$^{-/-}$ (n=4-10) rats. **(D)** TCRαβ$^{+}$CD4$^{+}$FoxP3$^{+}$ (CD4$^{+}$ Tregs) and TCRαβ$^{+}$CD4$^{-}$FoxP3$^{+}$ (CD8$^{+}$ Tregs) T cells and **(E)** TCRαβ$^{+}$CD4$^{+}$FoxP3$^{-}$ (CD4$^{+}$ Teffs) and TCRαβ$^{+}$CD4$^{-}$FoxP3$^{-}$ (CD8$^{+}$ Teff)s from total splenic cells from *Il34*$^{+/+}$ (n=4-9) and *Il34*$^{-/-}$ (n=4-10) rats were analyzed for expression of several markers before and after 3 days stimulation with anti-CD3 (1 ug/mL) and anti-CD28 (10 ug/mL) mAbs. **(F)** Regulatory TCRαβ$^{+}$CD4$^{+}$CD25$^{+}$CD127$^{low}$ (CD4$^{+}$ Tregs) and TCRαβ$^{+}$CD4$^{-}$CD45RC$^{low/-}$ (CD8$^{+}$ Tregs) T cells from *Il34*$^{+/+}$ (n=5-9) or *Il34*$^{-/-}$ (n=4-7) rats were sorted and tested for proliferation capacity in response to increasing concentration of anti-CD3 (0.25-0.5-1 ug/mL) and anti-CD28 (10 ug/mL) mAbs after 2 or 3 days of culture. Results are represented as mean ± SEM.

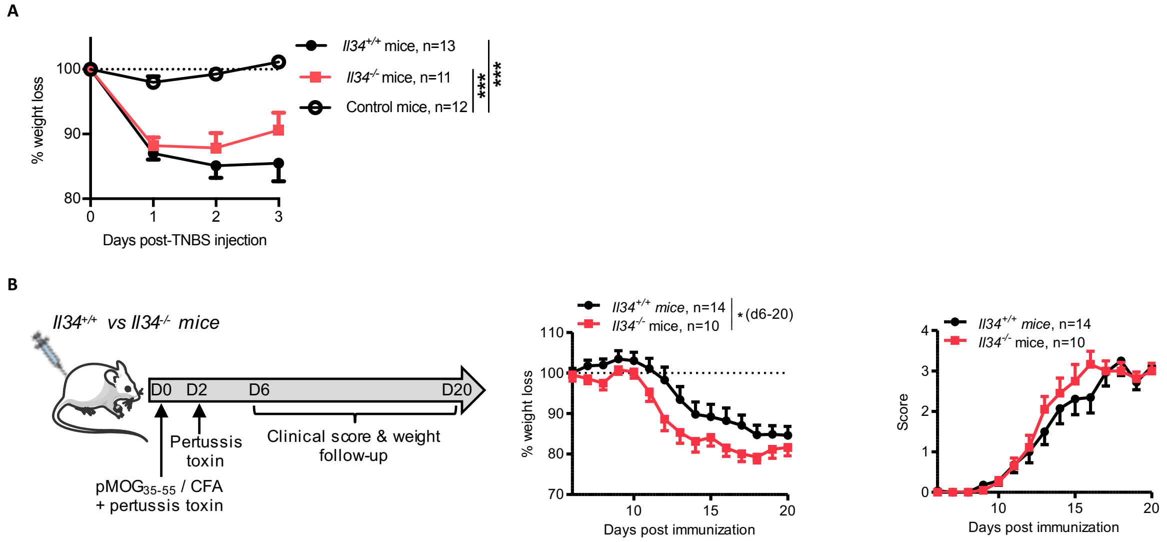


**Supplemental Figure 4. Colitis and EAE models in *Il34*$^{-/-}$ vs *Il34*$^{+/+}$ mice. (A)** *Il34*$^{+/+}$ and *Il34*$^{-/-}$ mice (n=11-13) were injected intrarectally with TNBS (100 mg/kg) in 50% ethanol. Control *Il34*$^{+/+}$ mice were injected with 50% ethanol alone. Mice weight loss was followed for 3 days. **(B)** Schematic showing the MOG-induced EAE model in mice. Adult *Il34*$^{+/+}$ (n=14) and *Il34*$^{-/-}$ (n=10) C57BL/6J mice of 8 weeks-old were immunized with the peptide $MOG_{35-55}$ and mycobacterium for EAE induction. Results are represented as mean ± SEM. Two-way ANOVA and a Bonferroni posttest for the weight loss and score analyses, * $p$<0.05, *** $p$<0.001.

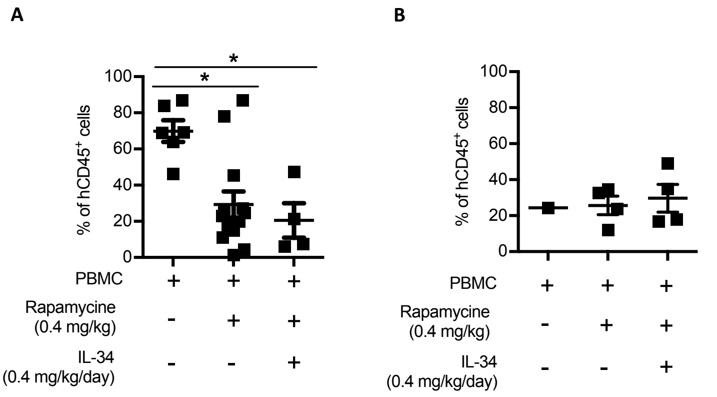


**Supplemental Figure 5. Human cell engraftment analysis in immune humanized mice**. At day 15 post-PBMC injection, FACS analysis was performed on human cells (hCD45) in the blood reflecting the engraftment (n=4-13) in the NSG GVHD **(A)** and skin graft rejection models **(B)**. Mann Whitney *U* test, * $p$<0.05.

Freuchet et al. Suppl. Figure 6

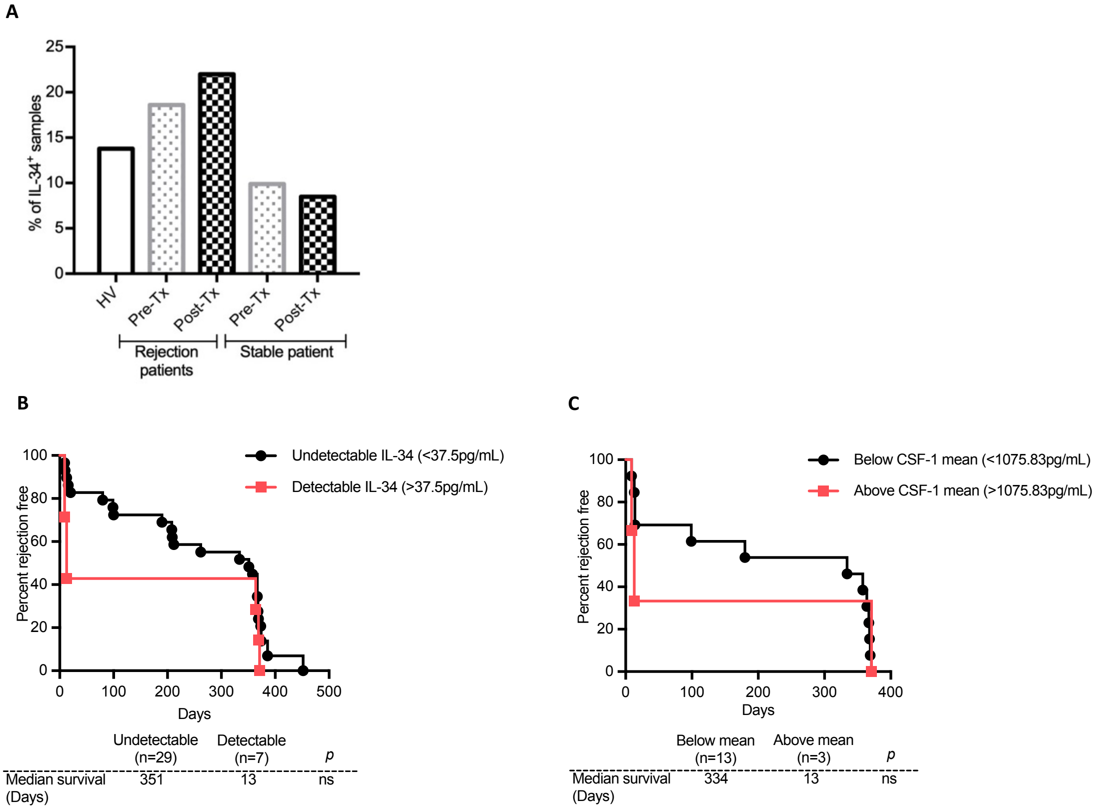


**Supplemental Figure 6. Correlation analysis of kidney transplant survival and IL-34/CSF-1 levels**. **(A)** The percentage of samples positive for IL-34 detectable levels was analyzed among the kidney transplanted cohort and healthy volunteers (HV). Graft survival of patients with an acute rejection episode occurrence according to IL-34 detectability post-Tx (undetectable: ≤ 37.5 vs detectable: ≥ 37.5 pg/mL) **(B)** or CSF-1 mean expression (below vs above 1075.83 pg/mL) **(C)**. Log- rank tests, ns *p*>0.05.

**Supplemental Table 1.** Antibodies used in this study

| Marker | Clone | Provider |
|---|---|---|
| hCD45 | HI30 | BD Biosciences |
| rTCRαβ | R7/3 | BD Biosciences |
| rCD4 | OX35 | BD Biosciences |
| rCD45RC | OX22 | Hybridoma from ECACC |
| rCD25 | OX39 | In-house |
| rCD127 | 717519 | Biotechne |
| rCD45RA | OX33 | BD Biosciences |
| rCD45R | His24 | BD Biosciences |
| rCD45 | OX1 + OX30 | BD Biosciences |
| rSIRPα | OX41 | In-house |
| rCD161 | 3.2.3 | In-house |
| rCD11b/c | OX42 | In-house |
| rCD103 | OX62 | In-house |
| Granulocytes | RP-1 | BD Biosciences |
| Granulocytes | HIS48 | BD Biosciences |
| rIgM | MARM-4 | In-house |
| rIgD | MARD-3 | In-house |
| rCD8 | OX8 | In-house |
| rCD44 | OX49 | BD Biosciences |
| rCD163 | ED2 | Bio-Rad |
| rMHC-II | OX6 | In-house |
| rFoxP3 | FJK16S | eBiosciences |
| rCD68 | ED1 | Bio-Rad |
| rCD71 | OX26 | In-house |
| rCD27 | LG.3A10 | In-house |
| rCD28 | JJ319 | In-house |
| rCD40L | AH.F5 | Biogen |
| rCD62L | OX85 | In-house |
| rICOS | JTT.1 | In-house |
| rCD122 | L316 | In-house |
| rIL-10 | A5-4 | BD Biosciences |
| rIFNγ | DB-1 | BD Biosciences |
| rCD3 | G4.18 | BD Biosciences |
| mCD3 | 500A2 | BD Biosciences |
| mCD4 | RM4-5 | BD Biosciences |
| mCD45RC | DNL1.9 | BD Biosciences |
| mFoxP3 | FJK16S | eBiosciences |
| mIL-34 | Polyclonal | Biotechne |

**Supplemental Table 2.** Description of groups and demographic characteristics of the cohort analyzed for IL-34 and CSF-1 serum levels.

| **IL-34** | | | | | |
|---|---|---|---|---|---|
| | **HV** | **Rejection patients** | | **Stable patients** | |
| | | **Pre-transplantation** | **Post-transplantation** | **Pre-transplantation** | **Post-transplantation** |
| *n* | 30 | 71 | 42 | 101 | 71 |
| *Age (y)* | 43.1±15.8 | 46.3±13.7 | 52±14.3 | 48.7±12.9 | 46.8±12.5 |
| *% of women* | 30% | 46.7% | 26.8% | 35.3% | 32.4% |
| *IL-34 Positive samples (%)* | 13.7% | 18.6% | 22% | 9.9% | 8.5% |
| *CNI* | / | / | 42/42 | / | 24/24 |
| *Tacrolimus* | / | / | 41/42 | / | 23/24 |
| **CSF-1** | | | | | |
| | **HV** | **Rejection patients** | | **Stable patients** | |
| | | **Pre-transplantation** | **Post-transplantation** | **Pre-transplantation** | **Post-transplantation** |
| *n* | 20 | 65 | 22 | 14 | 15 |
| *Age (y)* | 40.7±16.9 | 46.7±13.3 | 53.91±13.1 | 45±11.5 | 42.6±13.1 |
| *% of women* | 29.4% | 49.2% | 39.1% | 28.6% | 33.3% |
| *CNI* | / | / | 16/16 | / | 9/9 |
| *Tacrolimus* | / | / | 16/16 | / | 8/9 |
